# HPM Live μ for a full CLEM workflow

**DOI:** 10.1101/2020.09.03.281956

**Authors:** Xavier Heiligenstein, Marit de Beer, Jérôme Heiligenstein, Frédérique Eyraud, Laurent Manet, Fabrice Schmitt, Edwin Lamers, Joerg Lindenau, Mariska Kea-te Lindert, Jean Salamero, Graça Raposo, Nico Sommerdijk, Martin Belle, Anat Akiva

## Abstract

With the development of advanced imaging methods that took place in the last decade, the spatial correlation of microscopic and spectroscopic information - known as multimodal imaging or correlative microscopy (CM) - has become a broadly applied technique to explore biological and biomedical materials at different length scales. Among the many different combinations of techniques, Correlative Light and Electron Microscopy (CLEM) has become the flagship of this revolution.

Where light (mainly fluorescence) microscopy can be used directly for the live imaging of cells and tissues, for almost all applications, electron microscopy (EM) requires fixation of the biological materials. Although sample preparation for EM is traditionally done by chemical fixation and embedding in a resin, rapid cryogenic fixation (vitrification) has become a popular way to avoid the formation of artefacts related to the chemical fixation/embedding procedures. During vitrification, the water in the sample transforms into an amorphous ice, keeping the ultrastructure of the biological sample as close as possible to the native state. One immediate benefit of this cryo-arrest is the preservation of protein fluorescence, allowing multi-step multi-modal imaging techniques for CLEM.

To further explore the potential of cryo-fixation, we developed a high-pressure freezing (HPF) system that allows vitrification under different environmental parameters and applied it in different CLEM workflows. In this chapter, we introduce our novel HPF live μ instrument with a focus on its coupling to a light microscope. We elaborate on the optimization of sample preservation and the time needed to capture a biological event, going from live imaging to cryo-arrest using HPF. We will address the adaptation of HPF to novel correlation workflows related to the forthcoming transition from imaging 2D (cell monolayers) to imaging 3D samples (tissue) and the associated importance of homogeneous deep vitrification. Lastly, we will discuss the potential of our HPM within CLEM protocols especially for correlating live imaging using the Zeiss LSM900 with electron microscopy.

## 1. Introduction

Correlative Microscopy (CM) is an imaging approach that involves two or more complementary imaging techniques to obtain structural or molecular information of a given unique sample, usually by consecutive measurements. Here we will focus on one of the most popular correlative imaging techniques; correlative light and electron microscopy (CLEM). The conditions and the resolution at which the associated microscopes can work are directly determined by the physical properties of their imaging probes, i.e. light and electrons. Understanding the basic conditions required both for sample preparation and image optimization, is necessary when designing new CLEM project.

Light microscopy (LM) works at atmospheric pressure on hydrated materials, and therefore is the technique of choice for the dynamic imaging of specifically-labeled proteins. LM can visualize, with high throughput, large sample areas over hundreds of microns but only at a lateral resolution of ~200nm (not considering super-resolution techniques) and without contextual information.

In contrast, the high vacuum used in electron microscopy (EM) almost always necessitates dehydration and fixation of the biological samples. Hence EM yields static images in a relatively narrow field of view, from tens of microns to a few nanometers, with a relatively low throughput. However, EM can provide full contextual information down to sub-nanometer details, although specialized techniques (e.g. the Tokuyasu technique (Geuze, Slot, Van Der Ley, & Scheffer, 1981)) may be required to identify specific objects.

With a well-designed sample preparation/imaging workflow - where LM is typically used prior to EM - CLEM is able to preserve the benefits of the two techniques (de Boer, Hoogenboom, & Giepmans, 2015; Heiligenstein, Paul-Gilloteaux, Raposo, & Salamero, 2017; Müller-Reichert & Verkade, n.d.; T Müller-Reichert et al., 2014; Thomas Müller-Reichert & Verkade, 2012; Plitzko, Rigort, & Leis, 2009). This is particularly the case when high pressure freezing (HPF) is used, allowing direct cryofixation of biological samples without any chemical fixation.

Here we will demonstrate the use of the HPM Live-μ instrument as a flexible sample immobilization tool, allowing homogeneous vitrification of thick biological samples (up to 200 μm).

We will show the coupling of the HPF to an optical microscope minimizing the delay between dynamic imaging and cryoimmobilization, in order to fix the events of interest in both space and time.

## 2. High Pressure Freezing Principles

### i. Water phase diagram as a guide for HPF

With the increasing recognition of the importance of ultrastructure preservation during EM sample preparation, the use of cryo-fixation has become widespread. Plunge freezing vitrification (Dubochet, 2012) is the most broadly used approach because of its simplicity and low cost. However, due to heat transfer limitations, the sample thickness is restricted to a vitrification depth of less than one micrometer, significantly limiting the type of samples to be used in plunge freezing. In contrast, high pressure freezing (HPF) allows vitrification up to 200 μm deep (Moor, 1987; Shimoni & Müller, 1998; Studer, Michel, & Müller, 1989). Although HPF, compared to plunge freezing, requires more complex and more expensive instruments, it allows vitrification of a much wider variety of samples of biological material (e.g., cell culture, tissues) with close-to-native ultrastructure preservation.

At the basis of the need for HPF is the Leiden-Frost effect (Moor, 1987; Studer et al., 1989). When contacting a hot surface (the sample or the HPF chamber), liquid nitrogen (LN2) vaporizes and creates an insulating gas layer that reduces the cooling power of the system. This leads to slow cooling and hence to the ice crystallization in the sample, which can damage the ultrastructure and therefore hamper the interpretation of the observations. In HPF, the sample is exposed to a strong flow of LN2 through which, in a few milliseconds, the pressure surrounding the sample is elevated to 2100 bar while the temperature of the sample drops to 140 K (−133 °C, Fig. 1, P3), the gas being pushed through an outlet. Due to the high pressure and friction, the temperature of the liquid nitrogen during freezing will not drop to −196°C, but will reach −133°C, as shown in the water phase diagram (Fig. 1). It is important to note that during this rapid process the water does not reach thermodynamic equilibrium while traveling to its final temperature and pressure (P3). Under thermodynamic equilibrium and a constant pressure of 2000 bar, water solidifies below 251 K (−22°C) (Fig. 1, P2), as different ice forms can be present in at this point, the final ice formation will depend on other local variables. In the dynamic HPF process, however, where temperature and pressure are evolving simultaneously, water between 251 K and 176 K will go through a supercooled state (fragile liquid phase) before reaching the solid phase (Fig. 1, green line T_LL_). Below 176K the water will solidify into a “glass” (or strong liquid phase)(Tournier, 2020) that is composed of high density amorphous (HDA) and/or low density amorphous (LDA) ice (Tulk, Molaison, Makhluf, Manning, & Klug, 2019) depending on the exact pressure. Below 2000 bar, the majority will be LDA ice, while above 2000 bar, the majority will be HDA ice (Lin, Smith, Liu, Tse, & Yang, 2018). It is likely that water in HPF is a mixture of these two phases (Richter, 1994).

**Figure 1:**
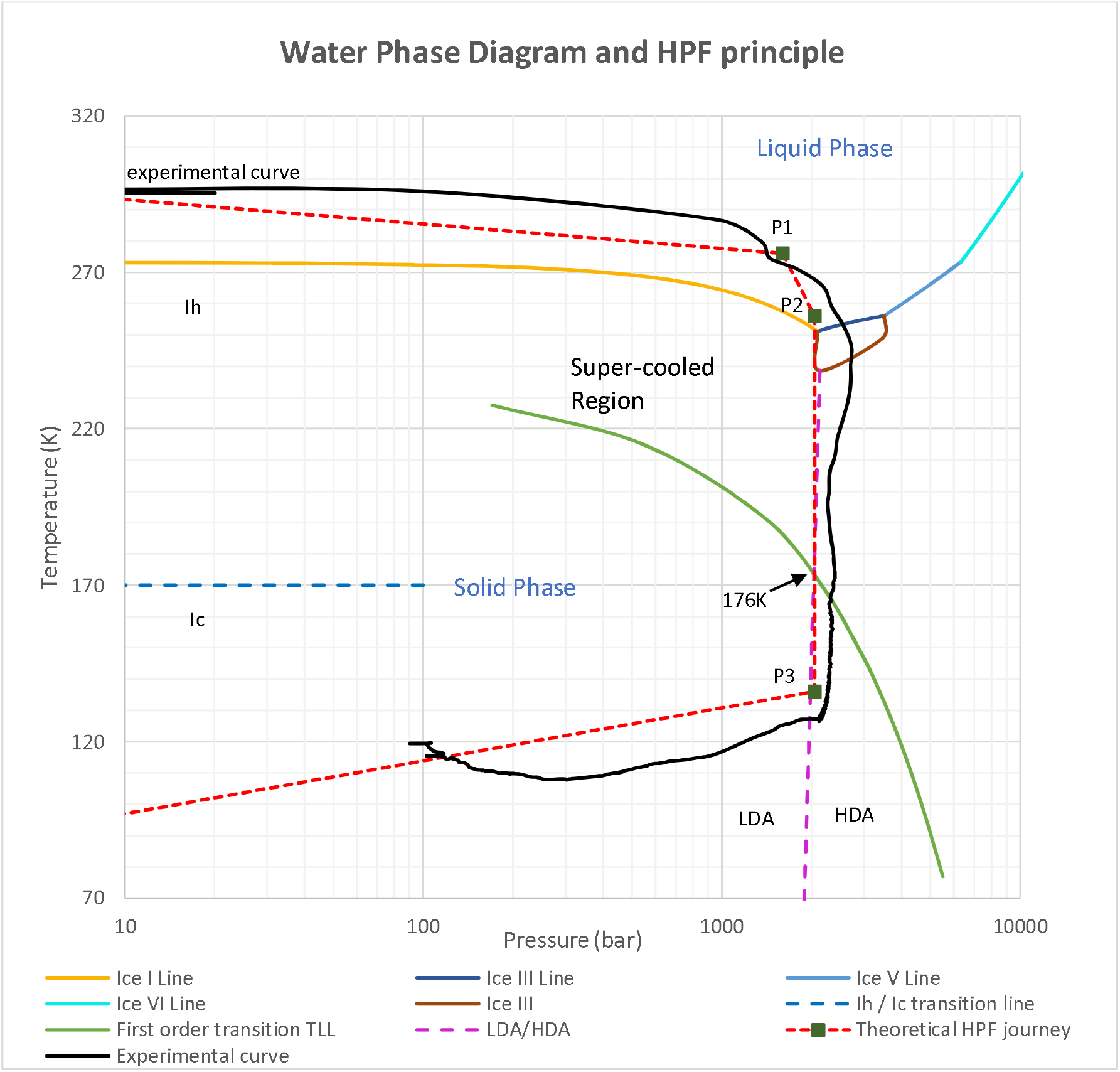
Water Phase Diagram in high pressure and a HPF experimental curve. Upon increasing the pressure, the freezing point progressively decreases until a minimum is reached at 2048bars and 251K (−22°C)(P2). Below the freezing point, and above the T_LL_ value (green line), water can achieve a supercooled state (unfrozen state) in a metastable state. Any perturbation in the equilibrium causes rapid crystalline transition (ice I line, I_h_ region). In High-Pressure Freezing, the conditions are raised to rapidly reach the 2048 bars equilibrium before fast cooling of the biological material to cross the super-cooled region and achieve water vitrification (a glass transition below T_LL_ where water does not crystalize). After vitrification, the system returns to atmospheric pressure while transferring the biological material in LN2 to prevent recrystallization in ice I_c_ near 138K (−135°C). The dashed red line represents approximately the journey expected to be imposed to the water molecules to achieve vitrification in HPF. The dark blue line (experimental curve) is one actual trajectory created by our HPM. The blue dotted one is the transition back from Ic to Ih.

### ii. The development of High Pressure freezing technology

HPF designs have been guided by the possibility to cross the phase diagram without ice crystal formation, traveling as quickly as possible along the line of the 2000 bar down to temperatures below 176 K. Hence, high pressure freezing requires synchronization of pressure build-up and sample cooling within only a small window of pressure and temperature to guarantee efficient vitrification. Historically, technologies to reliably generate a pressure of 2000 bar in less than 35 ms were exclusively hydraulic: oil pressure was accumulated and rapidly released to pressurize LN2 through a piston system. However, the pressure build-up remained relatively slow and synchronization required ethanol to slow down the sample cooling (McDonald, 2009; Shimoni & Müller, 1998). This widely acknowledged approach was commercialized in the HPM010 [ABRA Fluid AG, Widnau Switzerland] and the Wohlwend Compact 01 [Wohlwend GmbH, Sennwald, Switzerland]. To accelerate pressure building and optimize time correlation between pressure and temperature, Studer et al. developed the EMPACT technology [Leica Microsystems] that decoupled the pressurizing from the application of the cryogen (Studer, Graber, Al-Amoudi, & Eggli, 2001). Later on, Leica Microsystems developed an air pressure-based machine. Air pressure release is significantly faster than oil. Using a needle valve and LN2 channel conduction, this system improved the pressurization rate even though air pressure is less powerful due to the elasticity of air. The technology was commercialized as the Leica HPM100 [Leica Microsystems], and recently extended to apply light or electrical stimulation synchronized with the vitrification procedure (the EM ICE, Watanabe et al., 2011).

Another way to improve HPF vitrification, which is mainly important for the freezing of thick samples, is to increase the cooling rate, as was previously suggested by Shimoni and Müller (Shimoni & Müller, 1998). Using LN2 as the coolant, the cooling rate can be improved either by a higher flow of LN2, or by a faster contact, reducing the primary slow cooling caused by the nitrogen gas. Here, the challenge is to balance the pressure build-up and the sample cooling. A too rapid cooling will initiate water crystallization while too slow cooling will cause pressure damage on the sample.

## 3. Designing the HPM Live μ

### i. Improving the pressure rise speed and cooling rate

At 2000 bar, LN2 is at 113 K, compared to 77 K at 1 bar, which reduces the temperature gradient by 33K, and thereby also the cooling rate. To compensate for the strong effect on the temperature gradient, we significantly increased the LN2 entry speed and flow. In our ‘*Flash Core*’, we use proportional power technology (linear pressure increase) to control the hydraulic pressure which uniquely allows us to build pressure at a rate of 1000 bar/ms. With this proprietary technology we reach LN2 temperature and 2000 bar in only 2 ms, making our HPF system the fastest available up to date. We note that while the theoretical value for LN2 at 2000 bar is 113K, the LN2 working temperatures of HPMs in fact can reach 133 K, the difference being attributed to the high density of LN2 and friction during application of the high-speed LN2 jet (Shimoni & Müller, 1998; Stadie, Callini, Mauron, Borgschulte, & Züttel, 2015; Studer, Michel, Wohlwend, Hunziker, & Buschmann, 1995).

### ii. The HPM Live μ: A flexible research engine for optimized vitrification efficiency

Despite the long history of the use of HPF in many laboratories, most biological studies use the HPF instrument’s preset parameters: the machines are calibrated in the factory to reach a theoretical optimum and the numerical HPF values (time, temperature, pressure) are ignored until the EM observation when the quality of the specimen is revealed. At the end of a laborious sample preparation process, the user is left with returning to cryo-protection optimization, which could affect significantly the biological behavior of the sample. So far, no commercial system offered the capacity to tune the parameters that control the high-pressure freezing process.

All scientific approaches rely on the assumption that vitrification is simply a matter of rapid synchronized pressurization and cooling. It is our belief that the complexity of HPF is widely underestimated. As to what is progressively recognized in plunge freezing (Klebl et al., 2020; Shi, Ling, Zhu, & Zhang, 2019), HPF parameters should be tunable and accessible, in order to evaluate their impact on the sample (Leforestier, Richter, Livolant, & Dubochet, 1996). We have therefore designed the core of the HPM so that the user can set the key parameters of vitrification, including pressurization speed, cooling speed and cooling offset (pressure at 0°C), as well as the duration and value of the pressure applied.

As mentioned earlier, cryo-preservation is a kinetic process that occurs out of thermodynamic equilibrium (Tse, 2019; Tulk et al., 2019), and the dynamic parameters leading to the ideal HPF cooling curve still remains to be explored. With our ‘*Flash Core*’ technology, we are able to tune the rate at which LN2 is released under pressure. At the lower end, pressure is build up in 15 ms, comparable to other commercial HPF instruments. To widen the HPF control capabilities, we provide an ethanol injection option to shift the cooling peak to a later moment when the pressure is already established, achieving P/T synchronization delay of about 12 ms. The temperature delay for synchronization is partially due to the warming up of the ethanol in the chamber (25°C), which takes 7 ms. At the upper end, pressure builds up in just 2 ms, creating pressure before the cooling rate can trigger nucleation of the hexagonal ice.

Additionally, the ‘*Flash Core*’ technology allows the user to adjust the height and duration of the final pressure plateau, with which it becomes possible to explore different regions in the water phase diagram, e.g. to shift between HDA or LDA formation.

To ensure that the measurements are representing the conditions surrounding the sample, we placed the temperature and pressure sensors directly inside the HPF chamber, above and below the sample. Both variations in temperature and pressure are recorded at 1ms rate. This permits to explore the HPF parameters with accurate recording, and to adapt them to non-standard (e.g. thick, or non-aqueous) samples.

With the aim to broaden the user knowledge on the vitrification mechanisms of HPF, which remain empirical to date, most parameters (apart from the pressure rise speed) are adjustable through the graphical user interface. The user can adapt the HPF parameters to specific sample types (e.g., thickness, hydration) for better freezing results.

### iii. User interface for fast assessing and optimization HPF procedures

Vitrification by high-pressure freezing is a complex phenomenon, upstream of a delicate workflow, where either plastic embedded or cryogenic EM is conducted. As explained above, the quality of the sample is often validated only at the last step of the workflow: the EM analysis. When optimizing a protocol to improve sample preparation, it is critical to track the history of the sample: was fresh or fixed material used? Was cryo-protection used or not? Which HPF parameters were used, was postprocessing performed? To actively track sample quality and support data management imposed by journals for data reproducibility, all parameters associated to the vitrification step and related to the machine state are recorded in a standard ‘.csv’ file, simply accessible through the graphical user interface of the machine (Fig. 2, 3, 4). Optimization of protocols becomes therefore possible for each individual sample, allowing a more robust knowledge transfer. In addition, increasing metadata retention requirements for efficient traceability and reproducibility imply mandatory recording of HPF parameters and curve, and to give each sample a unique tracking identifier. This recording allows the tracking of each individual sample and allows the linking of light microscopy data with electron microscopy data in CLEM approaches. Therefore, the HPM Live-μ offers individual sample data recording and storage that can be related to its HPF parameters.

**Figure 2:**
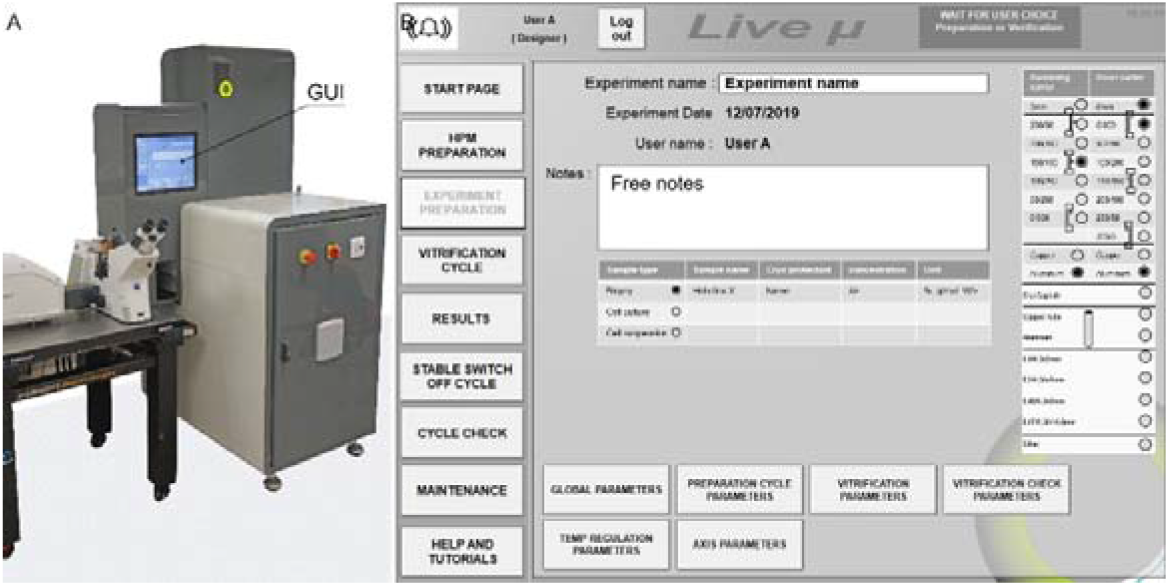
A system with a full Graphical User Interface (GUI). A-The machine displays full instructions to the user on a 15” touch screen (GUI). B-Each HPF shot gives rise to numerous measurements. All parameters are stored for later analysis. Sample’s metadata e.g., sample type, sample preparation such as pre-fixation or cryo-protection, carrier type etc. can be easily integrated in the note tab to track sample history

**Figure 3:**
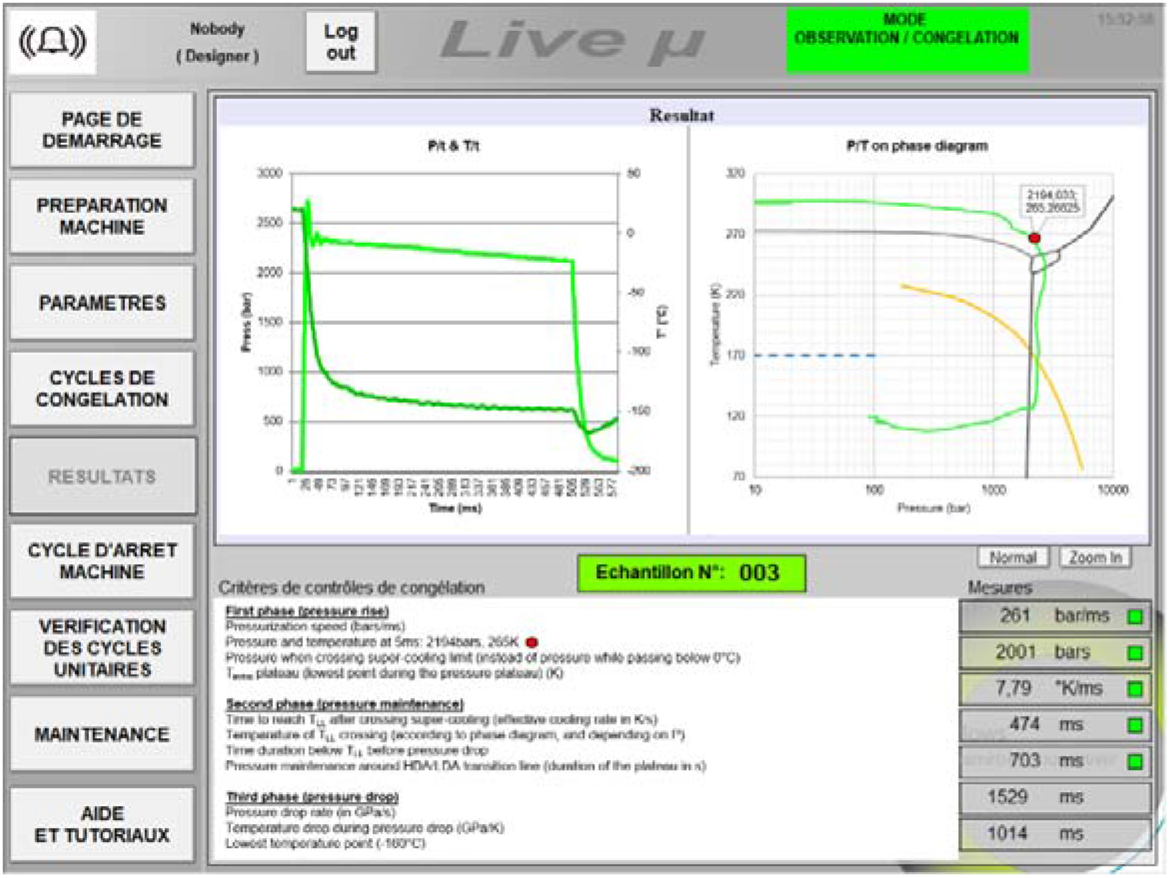
HPF shot analysis live to guide the user. After each shot, a series of parameters are measured and analyzed to guide the user in the quality control. This information can be retrieved for later comparative analysis (.csv format).

**Figure 4:**
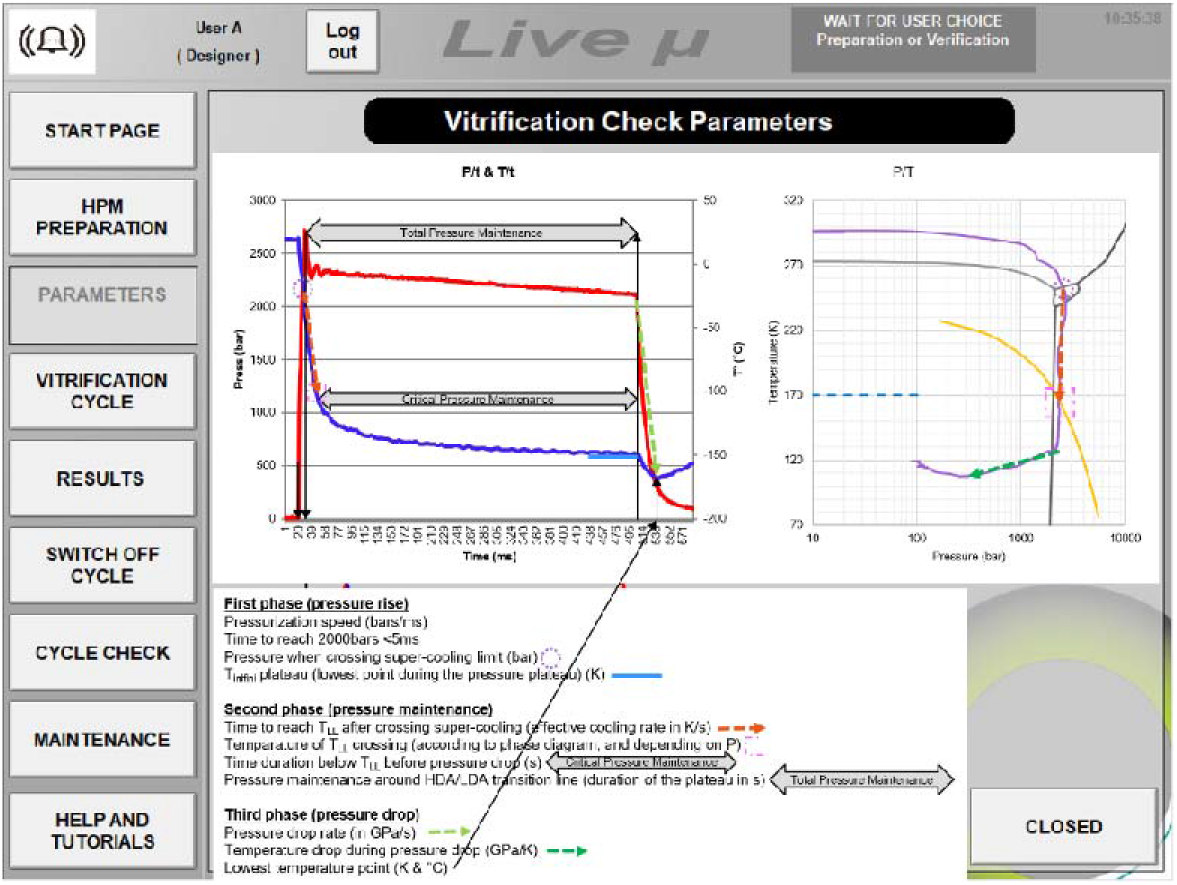
*Explanation of the measured parameters*. To help the user understanding the measured parameters, a page in the graphical user interface is available, describing in detail the measured parameters.

The presence of an Intel^®^ NUC mini-computer offers an advanced graphical user interface for a simple control of the system’s parameters and monitoring. The full HD chemical-proof touch-screen allows informative display of all the experiment’s parameters such as the temperature and pressure in the chamber (Fig. 3), for which also tutorial information is provided (Fig. 4). The ability to upload self-designed pdf tutorials supports the user’s autonomy and the need to adhere to facility’s management rules.

### iv. A wide range of sample holders for all commercial carriers

As high pressure freezing developed, various carriers were conceived in the field to match the experimental needs. The most common vessels that protect the sample during freezing are i) high pressure freezing carriers, ii) ‘tubes’ for cell suspensions, and iii) sapphire discs or ‘CryoCapsules’ for live CLEM (Fig. 5B). Below we will briefly discuss the differences between the vessels and give examples for high pressure freezing sample preparation for the different cases.

**Figure 5:**
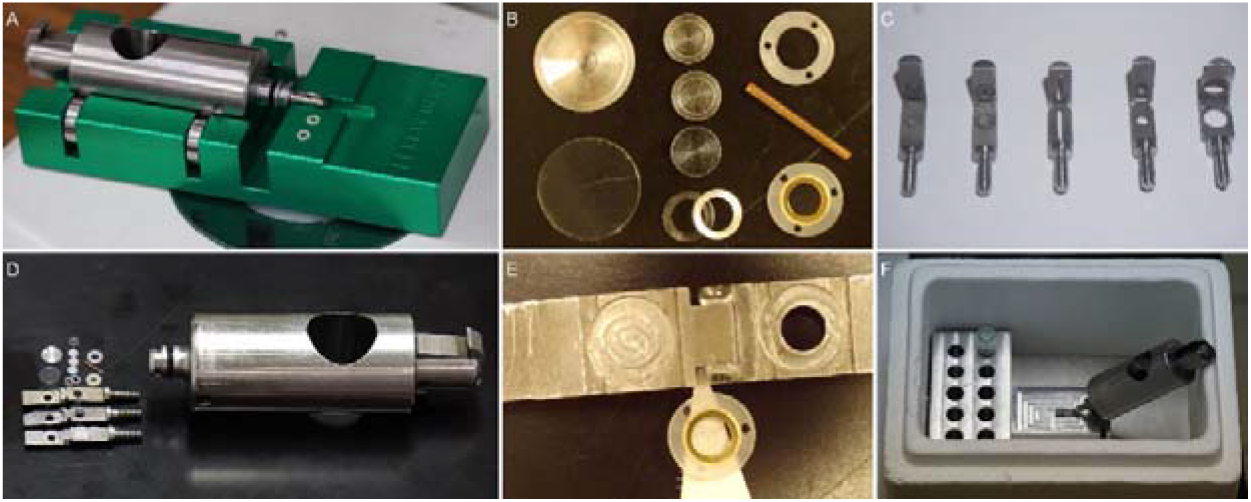
*Tools for efficient sample preparation and handling in HPF. A) loading station for the HPM Live μ., composed of sample holder (silver) and a stage (green)* B) Various existing carriers and sapphire discs for 3 and 6mm samples, cupper tube, and CryoCapsules. *C) Various stainless steel clamp to mount carriers adapted to the different sample* carriers. D) Ensemble of carriers, clamps and sample holders of the HPM Live μ F) CryoCapsule clamp and the CryoCapsule. G) Unloading station with Quick Freeze Substitution storage bloc.

The most common vessels are the **carriers,** also known as planchets or hats (Fig. 6a). These are small disks with an outer diameter of 3 or 6 mm, that can contain sample in liquid or solid form (e.g cell suspension or tissues). The carriers are made of metal (aluminum, gold-coated copper, copper, titanium) to combine high heat conduction and high mechanical resistance.

**Figure 6:**
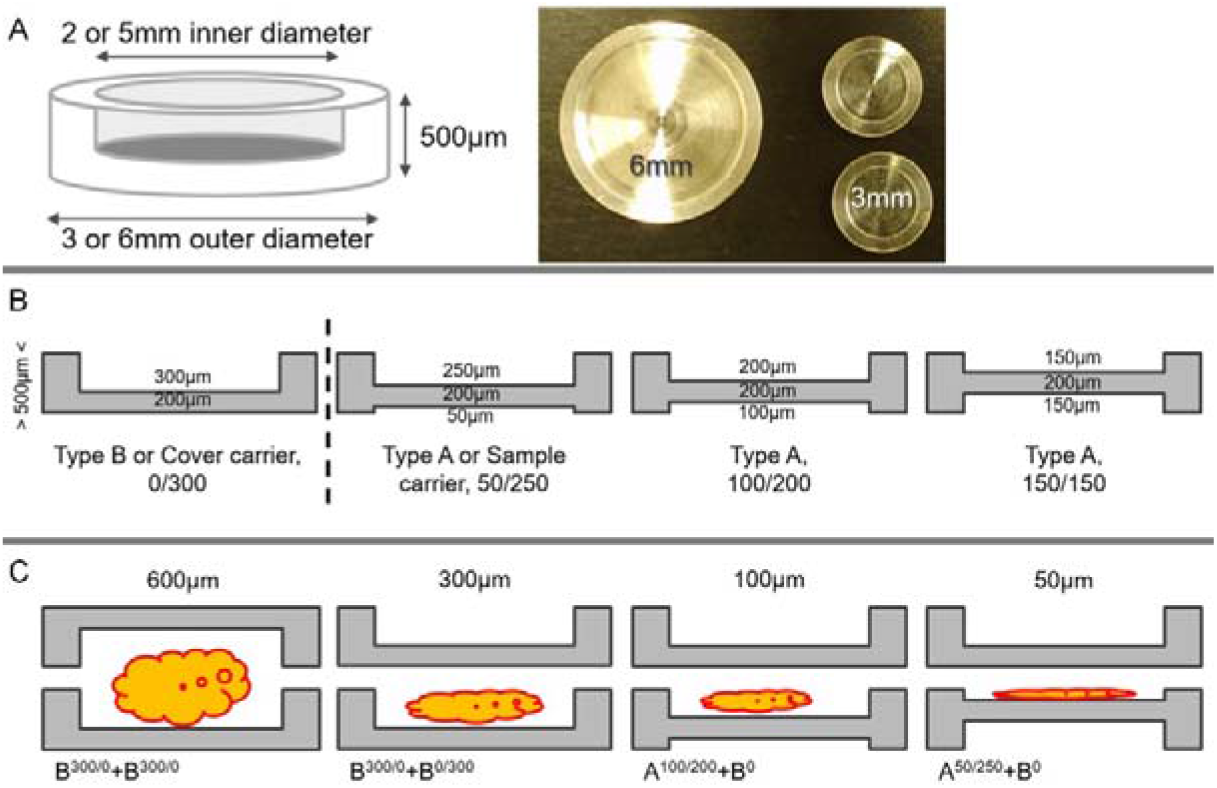
All carriers share common features. A- the outer diameter (3 or 6mm) defines the name of the carrier, the cavity has a diameter of 2 or 5mm accordingly and the total thickness is of 500μm. The assembly of 2 carriers will make 1mm to fill the clamp in a standard combination. B- the sum of the top and the bottom cavity always reaches 300μm. Type B carrier have a flat side and a cavity of 300μm. Type A carriers are selected to adjust the specific depth of the sample. C- the carriers assembly is chosen to fit tightly the thickness of the sample. To ease the orientation during sample manipulation, it is advised to deposit the sample in the cavity and cover it with the flat side of the type B carrier. All combinations can be used in accordance with the vitrification criteria.

A carrier (Fig. 6b) has a top and bottom cavity that together have a depth of 300 μm, and a central 200 μm thick support layer, and thereby a total height of 500 μm.

Although the most common carriers are of type A and B (for definition see fig. 6B), where A contains the cavity size of 100/200μm and B contains the cavity size of 300/0 μm (also named flat top), also other sizes of the carrier cavities are available, (25/275μm, 50/250 μm and 150/150μm).

To improve the freezing process it is important to choose the shallowest cavity possible to ensure the fit of the specimen, while reducing the vitrification volume to the minimum amount. Excess of water and increasing thickness will reduce the chances of homogeneous and high quality vitrification.

Before freezing, the samples are “sandwiched” between two carriers that fit the desirable thickness of the sample (Fig. 6c). Depending on the follow-up experiment after vitrification, the popular set-ups of the carriers will be: (Fig. 6c)

a. A bottom carrier with a cavity of any size and a flat carrier (type B) on top for follow up experiments such as cryo-FIB/SEM (Akiva et al., 2015, 2019), freeze substitution or cryosectioning
b. Two carriers with cavities of any size. This will be mainly used for freeze fracture, for example using cryo-SEM.

When ready, the sample is loaded into an appropriate clamp for HPF (Fig. 5c)

**Tubes** (Fig. 7A) are generally made of copper and are 9mm long. the outer diameter is 0.45, 0.5 or 0.6 mm and the inner diameter is 0.3, 0.35 and 0.3mm respectively.

**Figure 7:**
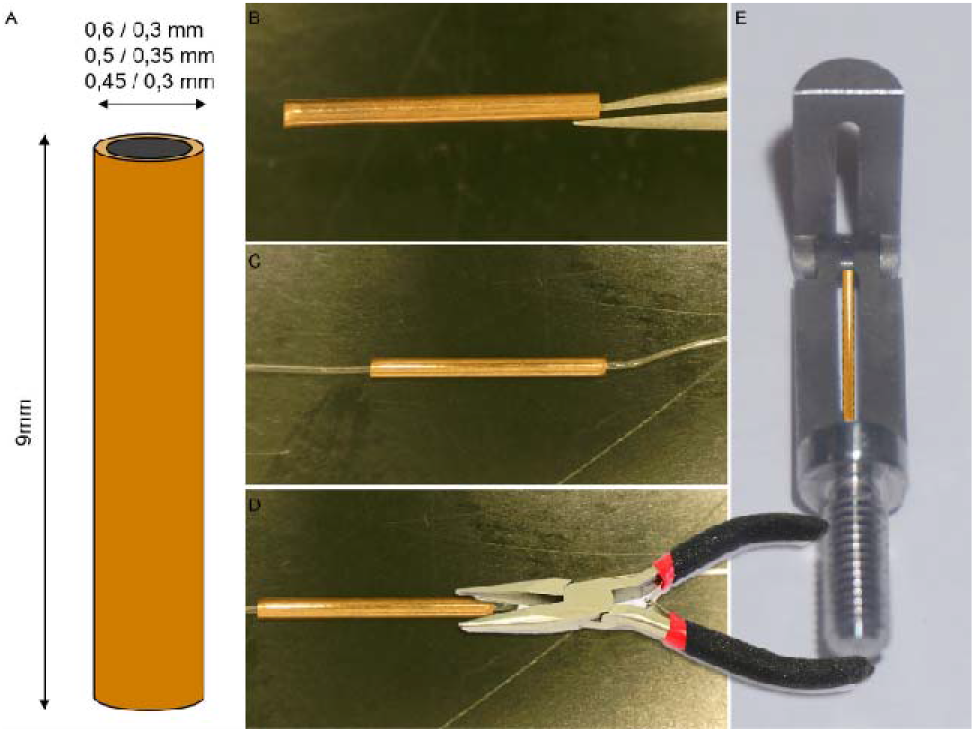
Cell suspensions freezing in a microcapillary tube. A- Schematic representation of the tube with the available dimensions. B- a 9mm long copper tube is used to protect the sample from the nitrogen jet during vitrification. C- insert the microcapillary tube in the copper tube and let the cell suspension fill the microcapillary. D- Seal both ends of the copper tube with pliers then cut off the outstanding microcapillary with a sharp razorblade. E- Deposit the copper tube in the clamp and proceed with freezing.

It is important to note that copper is highly cytotoxic and the cells suffer rapidly from a direct contact with it. Therefore, it is strongly recommended to load the cells in a capillary tube, itself loaded into the copper tube to protect the sample from the LN2 shot as described in the above section.

**Sapphire discs** (Monaghan, Cook, Hawes, Simpson, & Tomley, 2003), Aclar^®^ discs (Jiménez, Humbel, van Donselaar, Verkleij, & Burger, 2006) and CryoCapsules (Heiligenstein et al., 2014) are different freezable discs that enable the growth of cell monolayer on a transparent surface prior freezing. This approach is beneficial for CLEM studies. While both the Sapphire and Aclar^®^ discs are mounted between two metal carriers before HPF, the CryoCapsule is a self-standing system that enables the live imaging, seconds before freezing. We will elaborate on the CryoCapsule in section 4.

#### Choosing the right sample preparation method for HPF

Below we highlight four protocols for sample handling for HPF that cover most biological material. Protocols are given for (a) cell suspensions, (b) cell cultures and (c) tissues.

##### a. Cell suspension

Option 1:

1. Insert a microcapillary tube of approximately 2cm long inside an open copper tube (9mm).
2. Plunge the extremity of the microcapillary in the cell suspension until it is fully loaded.
3. Close both ends of the copper tube with a pair of pliers
4. Cut the microcapillary that comes out of the copper tube
5. Load the tube in the dedicated clamp (Fig 5c, clamp #3; Fig 7e)
6. Screw the clamp in the sample holder until full lock
7. Load the holder in the HPM Live μ
8. Initiate freezing cycle (foot paddle or touch screen)

Option 2: adapted from McDonald et al (McDonald et al., 2010), Figure 8:.

1. Concentrate the cell suspension by centrifugation and supernatant removal
2. Take a 200μm pipet tip and seal the narrow end of the cone with wax or by pressing on a hot surface
3. Pipet in the sealed tip, from the broad end of the cone, 100μL of the concentrated cell suspension.
4. Centrifuge gently the sealed tip to concentrate further the cell suspension at the end of the sealed tip.
5. Load the ‘filled tip’ on a pipet
6. Cut the sealed extremity
7. Deposit 5 to 10μL of the cell suspension in a A carrier (100 or 200μm deep)
8. Close the lid with the flat side of a B carrier
9. Load the assembly in the appropriate clamp (3 or 6mm in diameter, #4 or #5)
10. Screw the clamp in the sample holder until full lock
11. Load the holder in the HPM Live μ
12. Initiate freezing cycle (foot paddle or touch screen)

**Figure 8:**
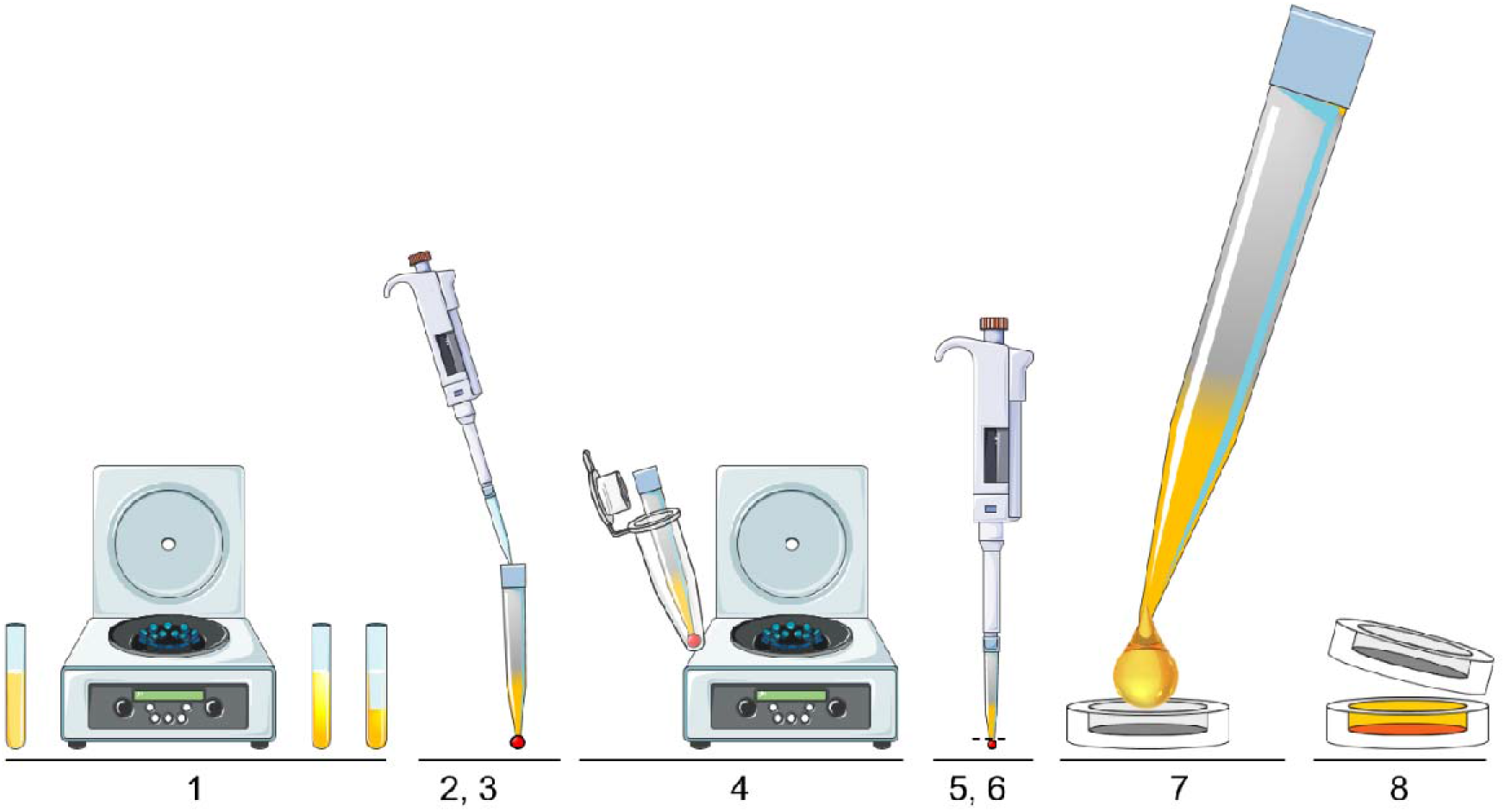
Preparation of cells suspension sample for freezing in a carrier. The steps 1 to 6 increase the cell’s concentration prior to loading in the carrier. The excess of liquid may reduce the freezing homogeneity. The final concentration (7, 8) is almost as high as in filtered cells.

##### b. Cell culture

There are many ways to prepare 2D cell culture for vitrification using HPF. Here we present five possible setup that are commonly used. Each method has its own advantage, and should be adjusted to the specific scientific question.

Choosing the support layer for cell culture (Fig. 9):

1. Free floating support such as sapphire (Reipert, Fischer, & Wiche, 2004) or Aclar^®^ disk (Jiménez et al., 2006). Requires a preliminary landmarking to identify the cells orientation throughout the sample manipulations (Culture, HPF, FS, Microtomy). Figure 9 B,C.
2. Free floating EM grid, precoated with formvar^®^, pioloform^®^ or a carbon film to support the cells at a sufficient density. Figure 9 D.
3. CryoCapsule, dedicated to Live-CLEM experiments, they comes ready to use out of the box (Heiligenstein et al., 2014). Figure 9 E.
4. Metal carrier with a differential depth (avoid carriers of type 150/150) to identify the cells orientation throughout the sample manipulations cells in the shallowest or deepest side accordingly. Figure 9 F Preparation for cell culture:
5. Sterilize the selected carrier by a brief wash in 70% ethanol followed by a quick rinse in PBS or distilled water.
6. Deposit the carrier in a petri dish, the carbon facing up to improve cell adhesion and growth (cell cultures tend to grow better on a thin carbon film). Figure 9 A.
7. Deposit the cell culture and let it settle for 2 days minimum and ensure appropriate adherence
8. Start your experimental cell culture protocol after a preliminary adherence of the cells.
9. On the day of the HPF day, renew the growth culture medium (the included carbohydrates act as a natural cryoprotectant). Preparation for freezing:
10. Pre-wet the carrier by dipping it in the cell culture solution (petri dish) to minimize the risk of bubbles
11. Load the disk covered with cells in a carrier (Fig 9b, c, d), cells facing the cavity
12. Close the lid with a wet cover carrier or the covering sapphire disc (CryoCapsule)
13. Load the assembly in the dedicated clamp (Fig 5C, clamp #2, 4,5)
14. Screw the clamp in the sample holder until full lock
15. Load the holder in the HPM Live μ
16. Initiate freezing cycle (foot paddle or touch screen

**Figure 9:**
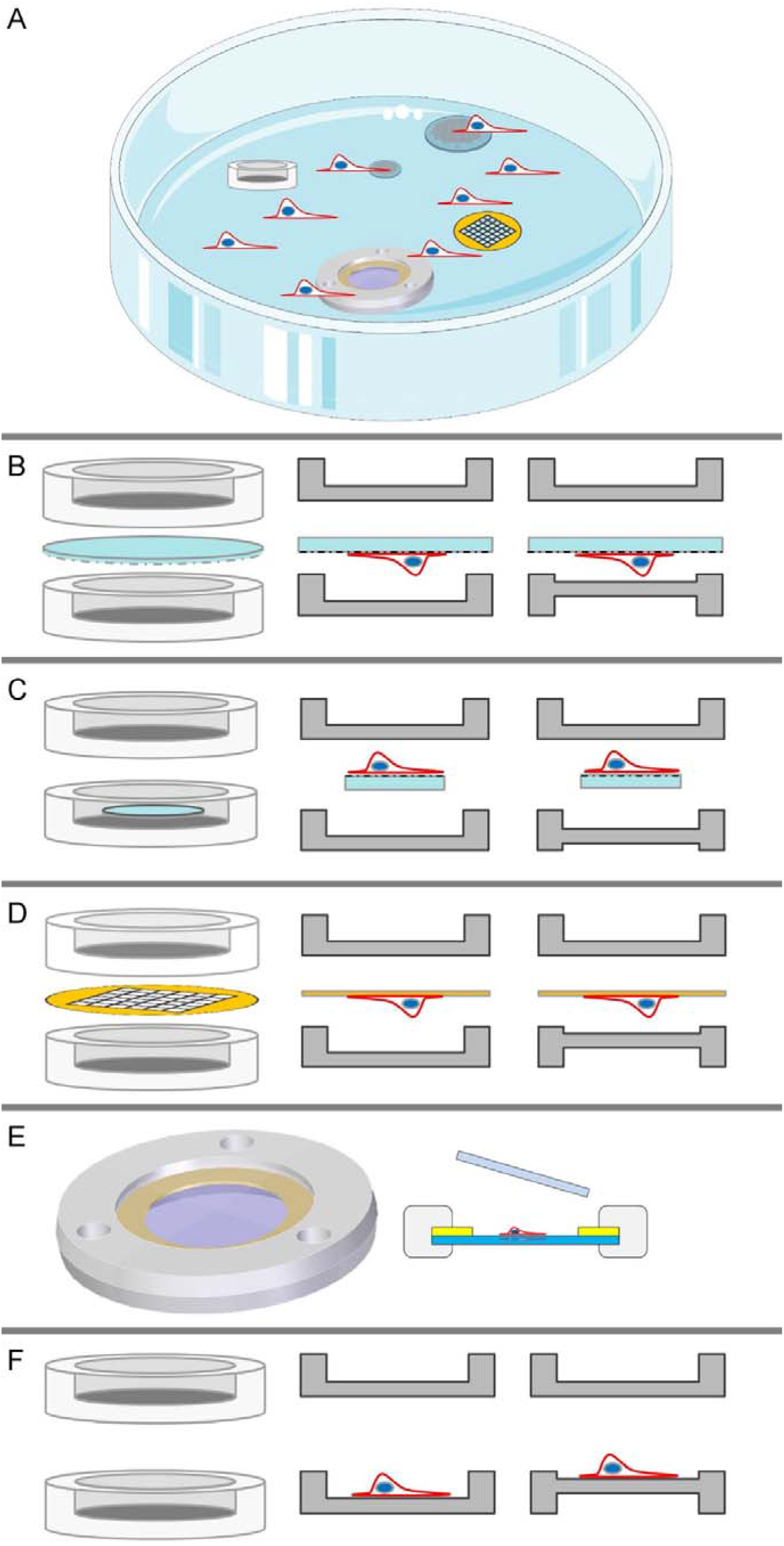
Cell culture growth options. A-all support described below can be cultured after sterilization in a regular petri dish, even simultaneously to figure out your best fitting protocol. B-a 3 or 6mm landmarked disc (sapphire or Aclar) is placed with the cells facing the cavity (chosen adequately for your experiment). A flat side type B carrier is used to complete the assembly in the clamp. The total thickness exceeds 1mm (disc extra thickness), so the clamp must be adapted (deeper or mildly flexible). C-the disc fits inside the cavity of the carrier, the cavity must be equal to the disk thickness plus the sample thickness. D-The cells are grown on top of a film coated EM grid, and the cells are oriented towards the cavity. The thickness of the grid is neglectable in the clamp assembly, not adaptation is required. E-the CryoCapsule is loaded in a specifically designed clamp with very tight measures to support live imaging and HPF without damaging the sapphire disc. F-the cells are grown directly inside the carrier. The depth of the carrier is chosen according to the sample thickness and closed with the flat side of the B carrier.

*Note: feed the cells with fresh medium one to two hours prior to your experiment in accordance to your protocol. Well-fed cells will vitrify better by up taking sugar present in the growth medium. For CLEM experiment consider to use medium without phenol-red to eliminate undesired fluorescence*.

##### c. Bulk specimen, tissue

Bulk specimen vitrification is often challenging: in most cases, the specimen is thicker than 200 μm, which is the vitrification depth limit. It is therefore advised to thin the sample prior freezing (vibratom, ultra-sharp razor blade, Feather^®^ scalpels) to the minimal thickness possible. Adapt the depth of the carriers to be the shallowest possible to lessen the water content and optimize freezing quality.

**Warning:** squeezing the sample with a too shallow assembly might impact the ultrastructure.

**Reduce the water amount** by using a filler (e.g., Yeast paste) or a cryo-protectant (CP) (10-20% Dextran 70000 KDa, BSA 10%). If possible, prepare the sample directly in the CP to impregnate it and once transferred in the carrier, top-up the carrier with the CP.

Ensure by **visual inspection that no air bubble is present:** air is compressible and will absorb part of the pressure, preventing the freezing ability of HPF. To ensure no air bubble will be created while closing, you can wet the flat side of the closing carrier with your CP solution before closure.

All combinations can be composed from 25 μm until 600 μm depth (figure 6 C). The shallowest will give the best results. Beyond 200 μm, it is strongly advised to use cryo-protectant for optimal preservation.

## 4. Integration of the HPM Live μ in a CLEM workflow

### i. Cryo-Capturing fast and small biological event

A key point for capturing dynamics in CLEM is the time difference between the last image collected during live imaging and the immobilization. In chemical fixation protocols, the specimen is usually immobilized by exposure to a fixative while imaging the sample progressively arresting (from milliseconds to minutes according to the biological specimen). The time correlation is 0 s if imaging is continued during the fixation (Stepanek & Pigino, 2017), but at the same time exposes the sample to the risk of introducing artifacts by the fixation process itself (Murk et al., 2003). An interesting alternative, that is yet to be commercialized is a microfluidic chamber that allows cryo-arrest during imaging (Fuest et al., 2018). Currently, no automated solution exists for CLEM of thick samples. The fastest procedures to date remain manual, preventing a close time correlation between the last image and the cryo-arrest (Heiligenstein et al., 2014; Verkade, 2008).

One way to overcome this constraint is to culture and observe the sample of interest in a HPF-compatible environment such as the CryoCapsule (Heiligenstein et al., 2014). The CryoCapsule is a cup-shapped vessel. The bottom is composed of a landmarked sapphire disc, and a 50μm thick spacer ring made of gold to create a cavity where the cells grow. The two elements are held together by a plastic wall with 4.5 mm external diameter and 2.8mm internal diameter. An independent covering sapphire lid fits inside the plastic walls to create an insulating environment for the cells during imaging and vitrification (Fig. 5 E, 6 A). The use of a sapphire bottom is currently the best compromise between mechanical resistance (Mohs: 9), thermal conductivity (35 W.m^-1^.K^-1^) and refractive index (1.76) (Heiligenstein et al., 2014). When a cell monolayer of interest is cultured in the CryoCapsule, the shortest light path is to observe the cells from below (Fig. 10 A). We therefore first optimized our setup to be compatible with inverted microscopes (Fig. 10 B).

**Figure 10:**
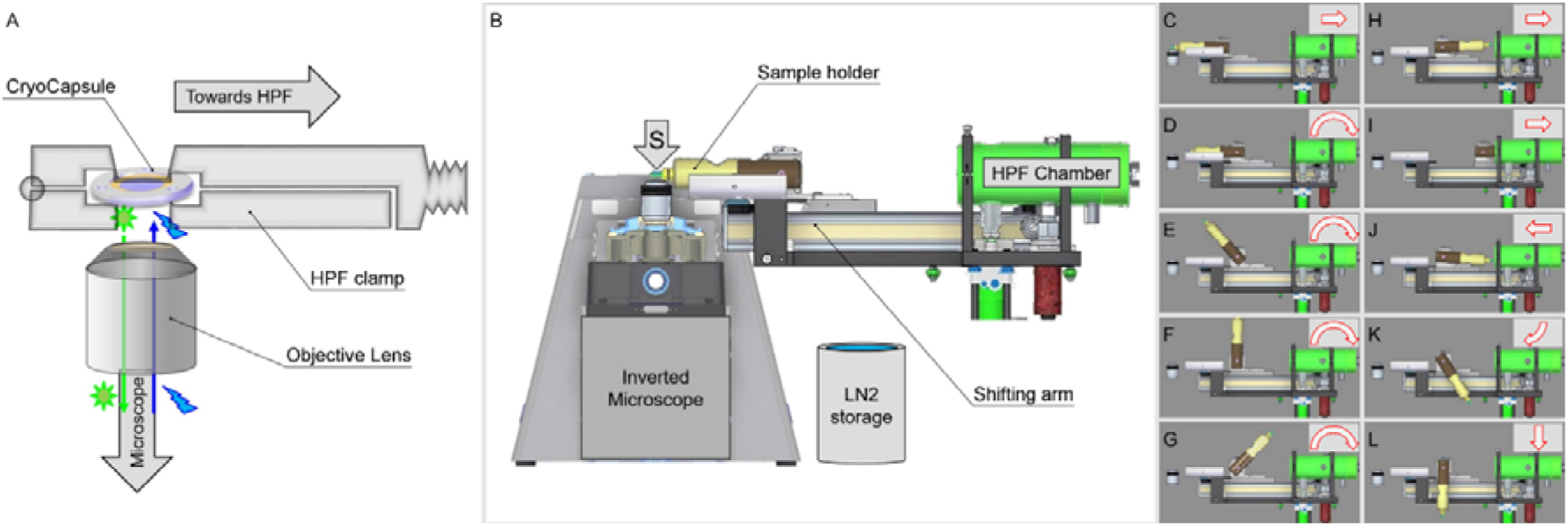
Physical path from live cell imaging to HPF. A) In CryoCapsule, the cells are lying at the bottom of the dish, in contact with the locating landmarks (carbon imprint), the shortest light path comes from the support sapphire disk underneath. B) The sample is resting at the tip of the sample holder, right above the microscope lens and close to the HPF chamber. C-L physical path of the sample between imaging(C), HPF (I) and storage in LN2 (L).

The best light microscopy set-ups use oil immersion lenses, maintaining a high numerical aperture of 1.4 N.A., which allows high sensitivity, shallow depth of focus and thereby a high resolution in all 3 dimensions. However, the poor heat conductivity of oil will reduce the vitrification capacity of any system and therefore must be removed before freezing. As our aim is to reduce the time window between the live imaging and the vitrification, it precludes the introduction of a blotting step between imaging and vitrification. Alternative lenses using water immersion (refractive index 1.333) would also require blotting as any remaining water would interfere with the vitrification of the observed material. Hence, the remaining solution is the use of air objectives for real-time CLEM followed by rapid freezing.

To address the points mentioned above and to enable fast transfer from live imaging to freezing, we designed a suspended arm, passively resting in imaging position (Fig. 10 B), waiting for the user to initiate the vitrification cycle. In practice, the freezing experiment follows these steps

1. Cell culture in a CryoCapsule (Fig. 9 E, 10 C)
2. Prepare the cells according to the defined experiment
3. Close the CryoCapsule with the covering sapphire disc directly in the culture petri dish
4. Blot away the excess medium
5. Load the CryoCapsule in the adapted clamp
6. Lock the clamp in the sample holder (live cell assembly)
7. Place the live cell assembly for imaging (Fig. 10 A,B)
8. Focus the microscope, locate the cell of interest
9. Initiate imaging (Fig. 10 C)
10. Once an event of interest is identified by the user, trigger vitrification (Microscope preset, HPM touchscreen or foot paddle)
11. The sample is automatically transferred into the vitrification chamber (Figure 10 C – I)

a. The sample retracts from the objective lens (step 10D)
b. Rotates 180° (step 10E-H)
c. Enters the HPF chamber (step 10 I)
d. Lock in secured position (Step 10 I)
e. HPF cycle starts. The sequence 10-11, occurs in 1.2 seconds, allowing the immediate freezing of many biological processes.
12. After vitrification, the sample is retracted (step 10J, K)
13. Direct release in a Dewar filled with liquid nitrogen (Steps 10 L)

The next steps in the CLEM experiment (storage, cryo-light microscopy, freeze substitution, (cryo-)FIB preparation) is according to the scientist’ project (Fig. 11).

**Figure 11:**
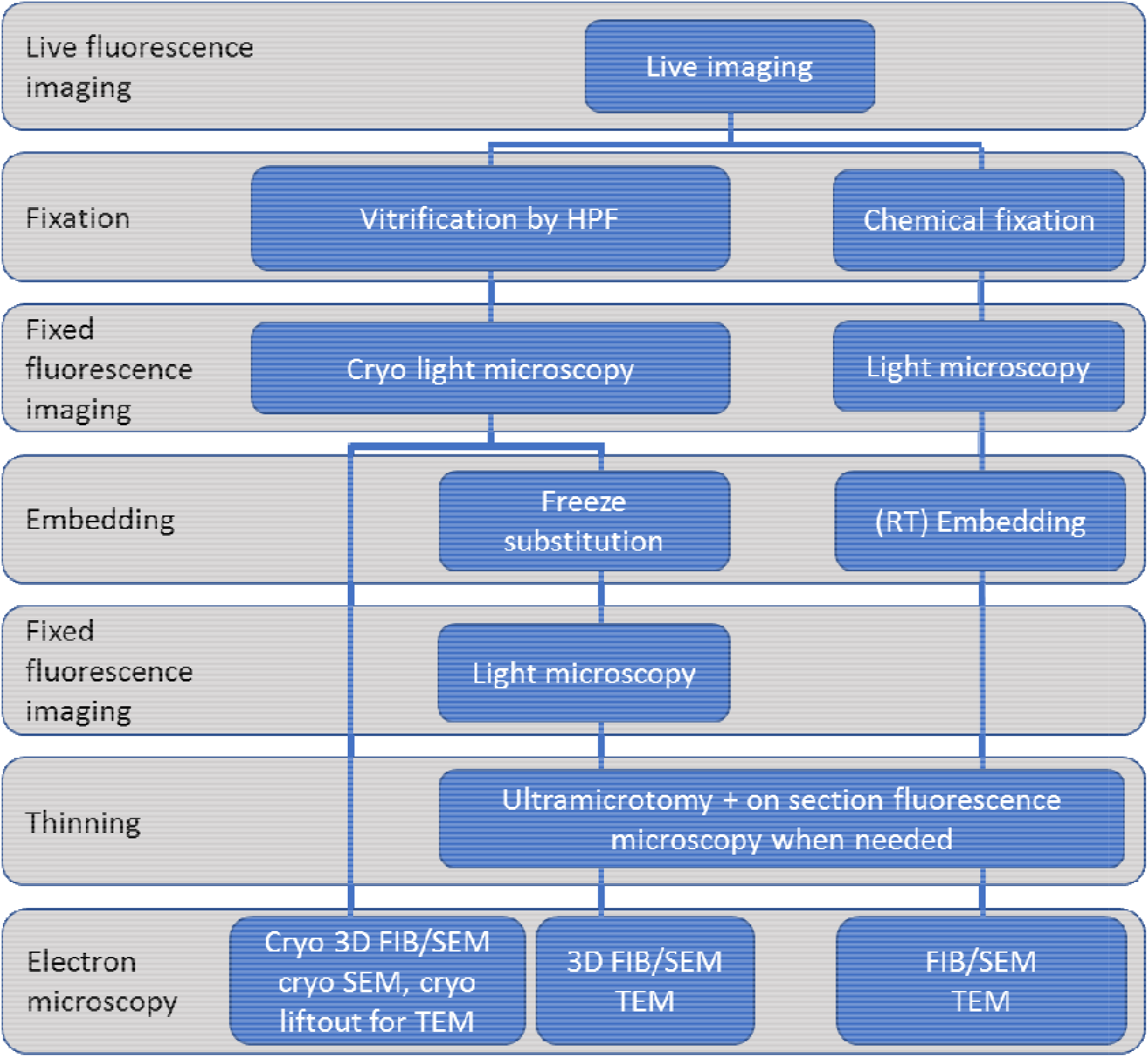
Simplified examples of workflows available from live fluorescence microscopy to electron microscopy. Each workflow shall be adapted to specific project and biological sample.

To maintain the CryoCapsule in position during live imaging and prevent damage during the blast of liquid nitrogen, the clamp was designed with specific shape and dimensions, to adequately direct the LN2 flow onto the sample during the vitrification. The depth positioning of the CryoCapsule defines the minimum working distance for the air-objective to be 1.8 mm (Fig. 10 A,B).

### ii. A flexible design to accommodate CLEM workflows

Correlative light and electron microscopy comes in as many flavors as do scientific projects. Typically, fluorescent light microscopy is conducted before any EM observation, as EM imaging usually involves heavy metal staining that quenches the fluorescence signal. Ideally, fluorescent imaging should be conducted at any step between live imaging and the final electron microscopy observation. A generic workflow adaptable to most scientific projects, in which the quality and fidelity is checked, could include the following steps (Fig. 11):

Although not all projects go through all these steps, we have designed our technology with this procedures in mind to make it adaptable to each scientific project and/or the main focus of the host facility.

**Note:** only three pathways are presented in the above scheme. Many diverted path can be used with expertise to accommodate the project’s complexity. Refer to your facility manager to explore a protocol that fits adequately your need.

In order to obtain the most direct light path possible for fluorescence imaging of living cells, the configuration of the HPF instrument has been designed to fit closely with inverted microscope bodies (Nikon TiE-2, Zeiss Axiovert), optimizing the signal detection of fluorescent proteins. If cryo-light microscopy is required, the microscope (mounted on rails) can be shifted away (Fig. 12 A,B), to fit an inverter arm and insert the cryo-objective into a cryostage. For cryo-light microscopy, the inverter arm is installed for live cell imaging on an upright microscope (Zeiss LSM 900, Fig 12 C, D). For cryo-fluorescence imaging, the cryo-arm is removed to fit natively the cryo-stage and optimize the most direct light path (Fig 12, E).

**Figure 12:**
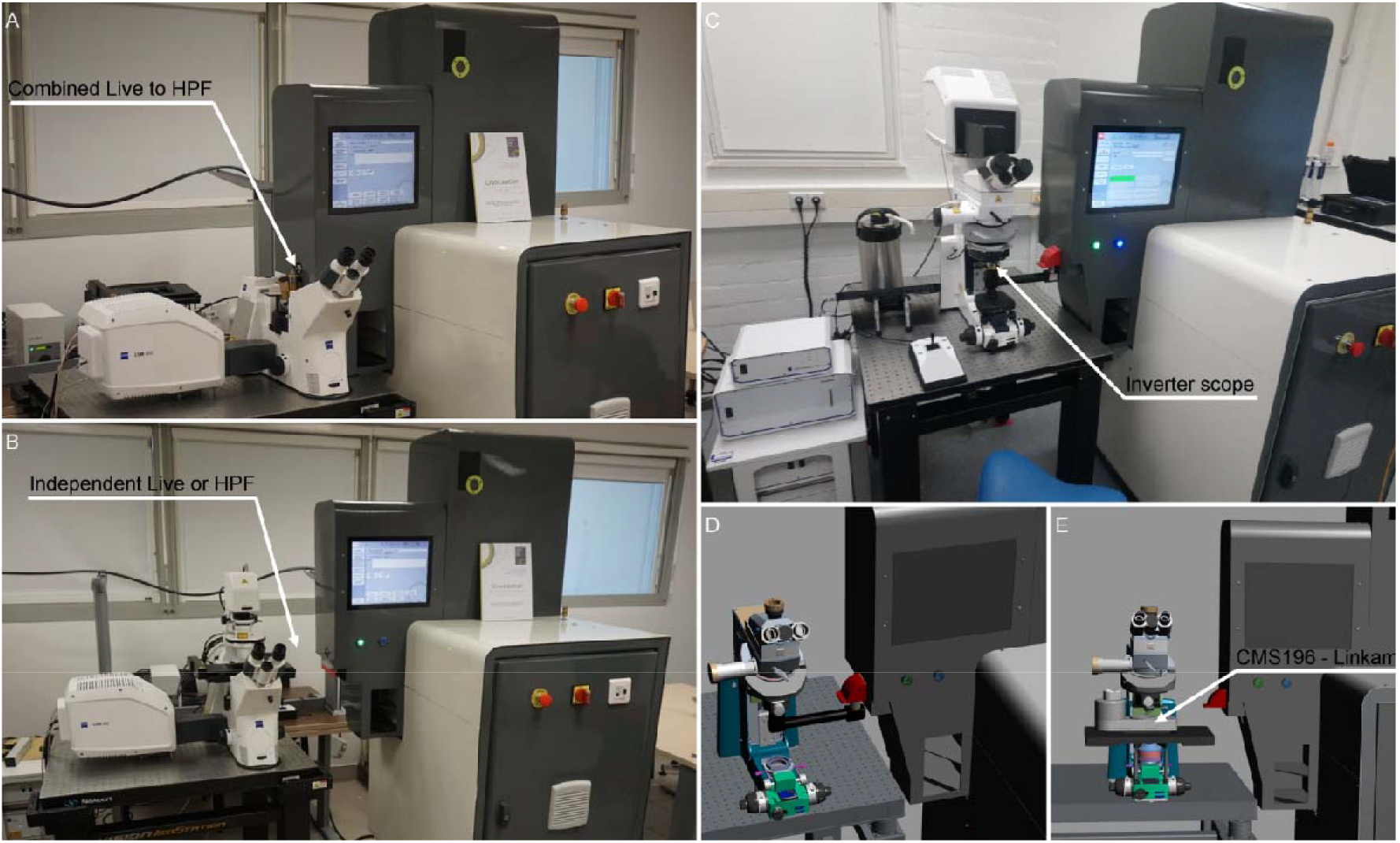
A flexible design to accommodate upright and inverted microscopes. A) The Axiovert Inverted microscope stage is tightly associated to the HPM Live μ. B) The system is separated to conduct independent live cell imaging, or cryo-light microscopy using an objective inverter. C-D) An upright microscope equipped with an inverter allows live cell imaging. E) The removal of the interver allows inserting a cryo-stage to achieve direct cryo-light microscopy.

In order to optimize equipment access, acknowledging that CLEM workflows are usually time consuming, the microscope can remain accessible for conventional LM use, decoupled from the HPM outside of HPF sessions.

## 5. A Live imaging, cryo-fluorescence, in-resin fluorescence and EM workflow: validation at each step

In CLEM, fluorescent live imaging enables the direct observation of a biochemical event, marked by fluorescent probes, where the wider biological context of the event can be observed with high resolution EM. By correlating the fluorescence and EM investigations, we could very precisely preselect the areas that we analyze in great detail with EM.

The use of EM is generally expensive, and often time consuming: not only the imaging and post processing, which can take hours (e.g. in 3D FIB/SEM), but also the sample preparation (metal staining, plastic embedding, ultra-microtomy). The ability to use light microscopy to pinpoint a specific event and validate specimen quality after freezing but before EM imaging is a revolution in correlative (3D) imaging, which saves hours of random exploring. Moreover, the possibility to check and compare the quality of a sample after live imaging, freezing, plastic embedding and EM observation will also limit the visualization of artifacts that were introduced during sample preparation. The benefits and possibilities of using light microscopy to guide EM imaging, and to validate sample integrity in CLEM, as will be discussed below, have so far been underemphasized.

Here we demonstrate how in three simple steps light microscopy can contribute to improving the CLEM workflow: by (i) evaluation of the HPF process under cryogenic conditions, (ii) assessment of the freeze substitution quality, and (iii) spatial correlation with EM. In this workflow, cells were grown directly on HPF carriers (25 to 100 μm deep) [Wohlwend GmbH, Sennwald, Switzerland]. When live-cell imaging was not possible, cells were first fixed with a mixture of polyformaldehyde (PFA) and glutaraldehyde (GA). It is important to note that although we will here discuss CLEM for room temperature, CLEM can also be performed under cryogenic conditions (cryoCLEM), limiting - if not preventing - the generation of sample preparation artifacts.

### i. Sample evaluation under cryogenic conditions after high pressure freezing

A common approach to the use of CLEM in cell studies is by growing 2D cell layers on TEM grids, and then to rapidly freeze these by plunge freezing (Dobro, Melanson, Jensen, & McDowall, 2010; Hampton et al., 2017; Kasas, Dumas, Dietler, Catsicas, & Adrian, 2003; Ravelli et al., 2020; Schultz, 1988) or high pressure freezing (Dahl & Staehelin, 1989; McDonald, 2009; Studer et al., 2001; Verkade, 2008). As the thickness of a cells generally exceeds 5 μm, HPF is the method of choice for vitrification, to prevent the formation of ice crystals that can damage the delicate cell structure.

HPF carriers can also conveniently be used for 3D cell cultures, creating a microenvironment for the cells. The depth of the carriers can be chosen, ranging from 25 μm to 300 μm. This also applies to the carriers in which one of the surfaces is a sapphire disc (as in the CryoCapsule), with the big benefit that these allow the cryocapture within 1.2 seconds from the live observation (Fig. 10), which is not possible with the metal carriers.

In the experiment depicted below (figure 13, 14), the steps were as follow

1. The cells are cultured on the carrier
2. Live fluorescent imaging of the cells prior freezing at intermediate magnification (20x) to create an overview (Fig. 13A)
3. High resolution imaging (60x airyscan mode) is acquired at specific location of interests (Fig. 13 A) Note: Depending on specific needs, samples may be embedded in a cryo-protectant solution (e.g., dextran 10-20%).
4. If the carrier is closed by a flat top cover (Fig 13 B) that must be removed after freezing, the flat side of type B carrier should be coated with a lipid film (e.g. phosphatidylcholine from egg yolk, diluted in ethanol to get 1% solution (wt/vol) (Studer et al., 2014)
5. After the sample is sandwiched between the two carriers, it is transferred to the HPF sample holder for freezing (Fig. 13 C, D).
6. the sample is high pressure frozen (Fig. 13 E, F)
7. the top carrier surface is removed (this step is facilitated by the lipid layer, step 4)
8. the sample is introduced into a cryo-stage that enables 3D imaging under cryogenic conditions in an environment that prevents ice contamination from the atmosphere Fig 13 G).
9. The stage is equipped with a designated holder for either TEM grids or HPF carriers.
10. Cryo-fluorescence imaging enables the first step in the validation process in CLEM sample preparation (Fig. 14). As most fluorescent probes are visible under the cryogenic conditions, this allows pre- and post-HPF imaging, to assess the effect of HPF, e.g. on the morphology of the cells, as well as on ice-formation.
11. The cellular morphology is evaluated using the fluorescent markers, as shown for fibrosarcoma cells (Fig. 14 F).

**Figure 13:**
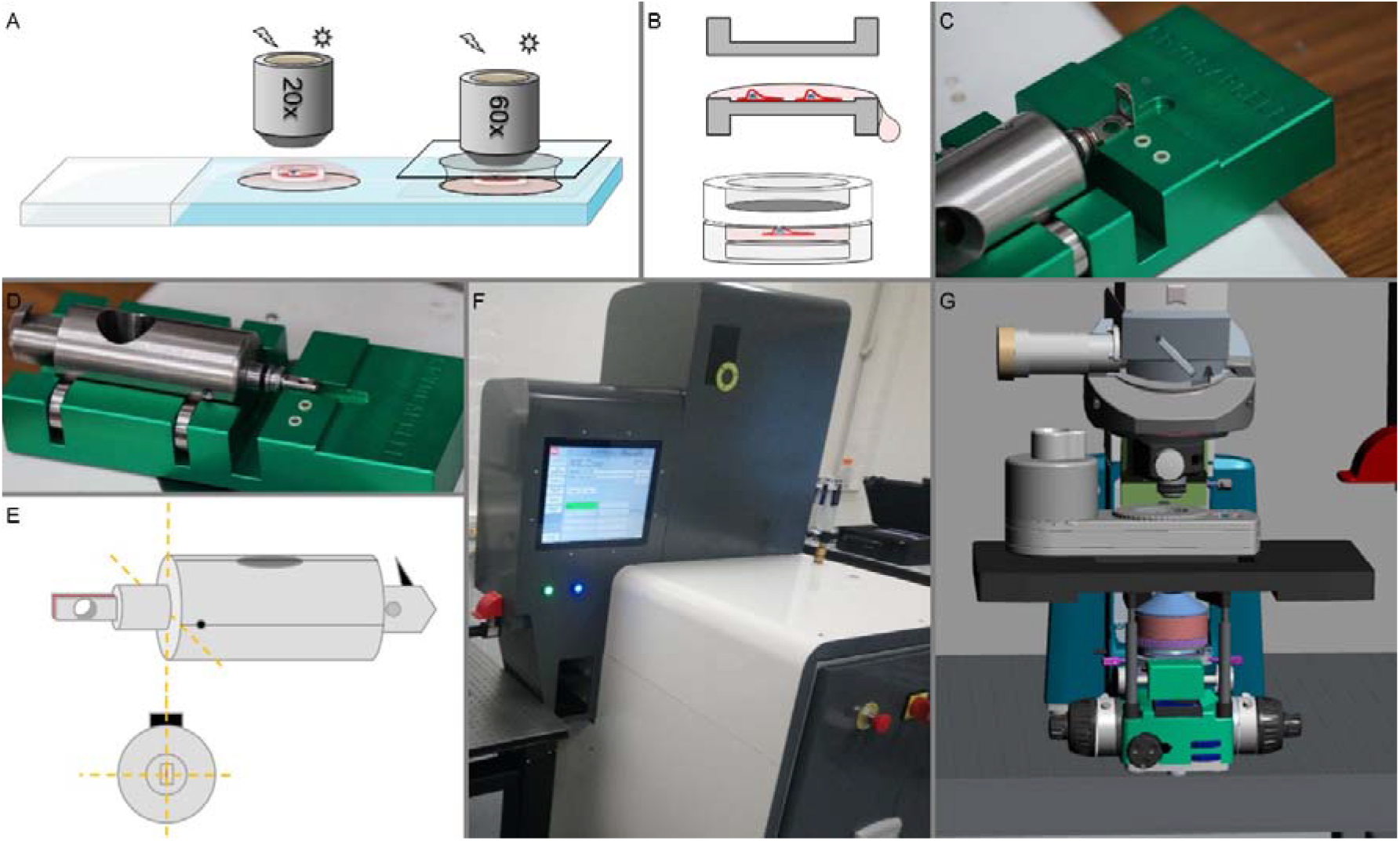
Live to CryoLM workflow. A-Cells grown directly on HPF carrier are imaged live in a drop of media on an upright microscope. B-The sample is enclosed using the flat side of a Type B carrier. The excess of medium is blotted away. C-The assembly, blotting and loading are done on the loading station. D-The clamp containing the carriers is screwed until full lock. E-The clamp is positioned vertically to ensure direct contact with the liquid nitrogen during the vitrification process. F-the sample is loaded into the HPM Live μ and vitrification is initiated. G-The sample is imaged at cryogenic temperature on a Linkam CMS196 stage prior to electron microscopy preparation.

**Figure 14:**
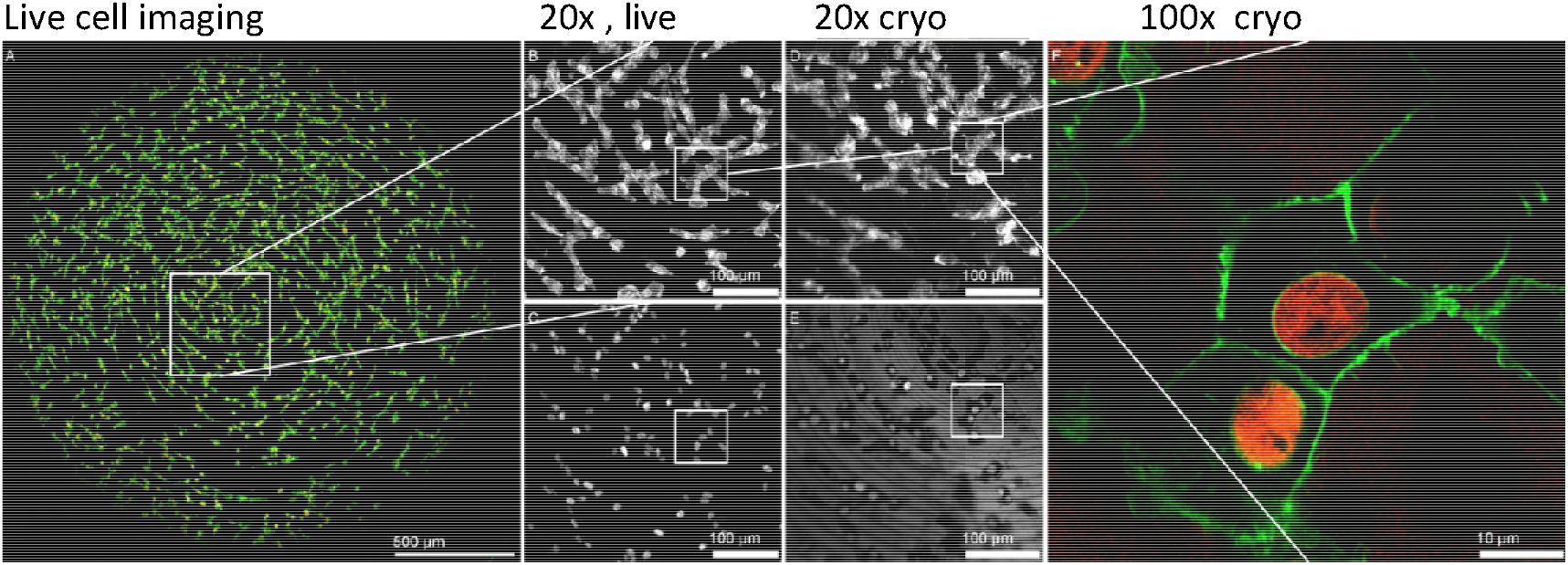
From live-cell imaging to cryo-fluorescence, post-high pressure freezing. Fibrosarcoma cells (HT1080; Histone 2B-mCherry and GFP-Lamin A) were grown directly on the HPF carriers without additional coating before seeding. After an addition plasma membrane staining (Cell mask green, Invitrogen) the live-cell images were taking using the Airyscan SR-4Y mode on an upright Zeiss LSM900 (A-C). (A) Overview of the complete carrier. The zoomed in area shows in Lamin A and plasma membrane after 488nm excitation (B) and Histone 2B after 561nm excitation (C). After HPF, images were taken under cryogenic conditions supplying the Zeiss LSM900 with a Linkam CMS196 stage (D-F). Same area as before freezing showed the preserved fluorescence in both 488nm excitation (D) and 561nm excitation (E). Note the in 561nm excitation there is increased reflection caused by the carrier. Finally, high resolution images (Pixel size 79×79×450 nm) can be obtained of area of interests (F) using a 100x long distance objective.

**Note:** after freezing, some cells appear to be swollen (Fig. 14). It is currently unknown if this is a physical artifact or the result of a change in the optical pathway, e.g. related to a change in refractive index. However, this apparent swelling disappears after freeze-substitution.

Interestingly, using a 561 nm excitation laser to image the sample in cryogenic conditions, a stronger background signal is detected due to a higher reflection of the carrier compared to live-cell imaging (Fig 14E). Although this is a disadvantage for cryo-fluorescence imaging, the fact that the carrier pattern is also seen in EM allows us to use the background features for navigation, to improve the correlation between both imaging modalities.

Of particular interest for possible cryo-EM imaging is that ice-formation can be easily assessed in cryo-fluorescence.

Using a cryo-protectant (e.g. 10% dextran) the samples appear more transparent, less milky when imaged in cryo-fluorescence. The milky appearance most likely relates to the formation of small (100-200 nm) ice crystals that are too small to be observed, but large enough to cause scattering. Hence, already in this early stage researchers can assess sample quality and make selections on which specimens to continue with for cryo-EM or freeze substitution.

### ii. Fluorescent preservation after freeze substitution

Freeze substitution (FS) is a popular method to prepare biological samples for EM due to its high degree of preservation of the tissue. During FS, the high pressure frozen sample is gradually brought up to room temperature while being exposed to contrast agents (heavy metal stains) and embedded in a resin (FEDER & SIDMAN, 1958; Walther & Ziegler, 2002).

FS is a time-consuming process that takes several days. The following electron microscopy procedures are even longer and more expensive. The ability to select the high quality samples after HPF-FS becomes therefore important to avoid inadequate electron microscopy processes.

Although, the heavy metal stains used in most protocols quench the fluorescence of the fluorescent probes, some protocols enable the preservation of fluorescence after FS (Johnson et al., 2015; Kukulski et al., 2011; Nixon et al., 2009; Peddie et al., 2014, 2017). Here we used the following process to evaluate the sample preservation prior to electron microscopy:

1. Cells were grown in aluminum carrier (figure 9F and 14)
2. Live cell imaging mapping (20x) is acquired
3. High Resolution detailed acquisition (60x, Airyscan mode) is acquired
4. HPF
5. FS in 0.2% Uranyl Acetate, 5%H2O in dry acetone, 5 hours, at −90°C.
6. Progressive warming (+5°C/hour) until −45°C
7. 3 rinses in dry acetone
8. 25% R221 impregnation for 2 hours
9. 50% R221 in acetone for 2 hours
10. 75% R221 impregnation in acetone for 2 hours and progressive warming to −30°C
11. 100% R221 impregnation overnight
12. 100% R221 impregnation in fresh solution and UV curing 48hours at −30°C.
13. Progressive warming +5°C/hour until 20°C
14. Collection of the sample
15. In resin fluorescence mapping (20x)
16. High resolution acquisition (60X Airyscan mode)

Note: the R221 resin (CryoCapCell, France), generates a low background and high contrast in scanning electron microscopy, and thus allows the use of a low concentration of uranyl acetate (0.2%) as a staining agent. Low uranyl concentration limits the quenching of the fluorescent probes present in the biological specimen. The R221 resin is compatible with standard fluorophores like AlexaFluor dyes, or fluorescent proteins such as GFP and mCherry.

Note 2: fluorescent probes which lost their activity under cryogenic conditions (e.g. DAPI and DsRed), were not reactivated after the FS resin (Kaufmann, Hagen, & Grünewald, 2014), while other probes which are visible both under cryogenic conditions and room temperature can be used (e.g DAPI can be replaced by Hoechst for nuclear staining)(Faoro et al., 2018; Heiligenstein et al., 2014).

The ability to preserve the fluorescent probes after embedding in resin, permits fluorescent imaging to assess the structural preservation of the samples after bringing them back to room temperature (Fig. 15), and to specifically determine the region of interest before moving to EM.

**Figure 15:**
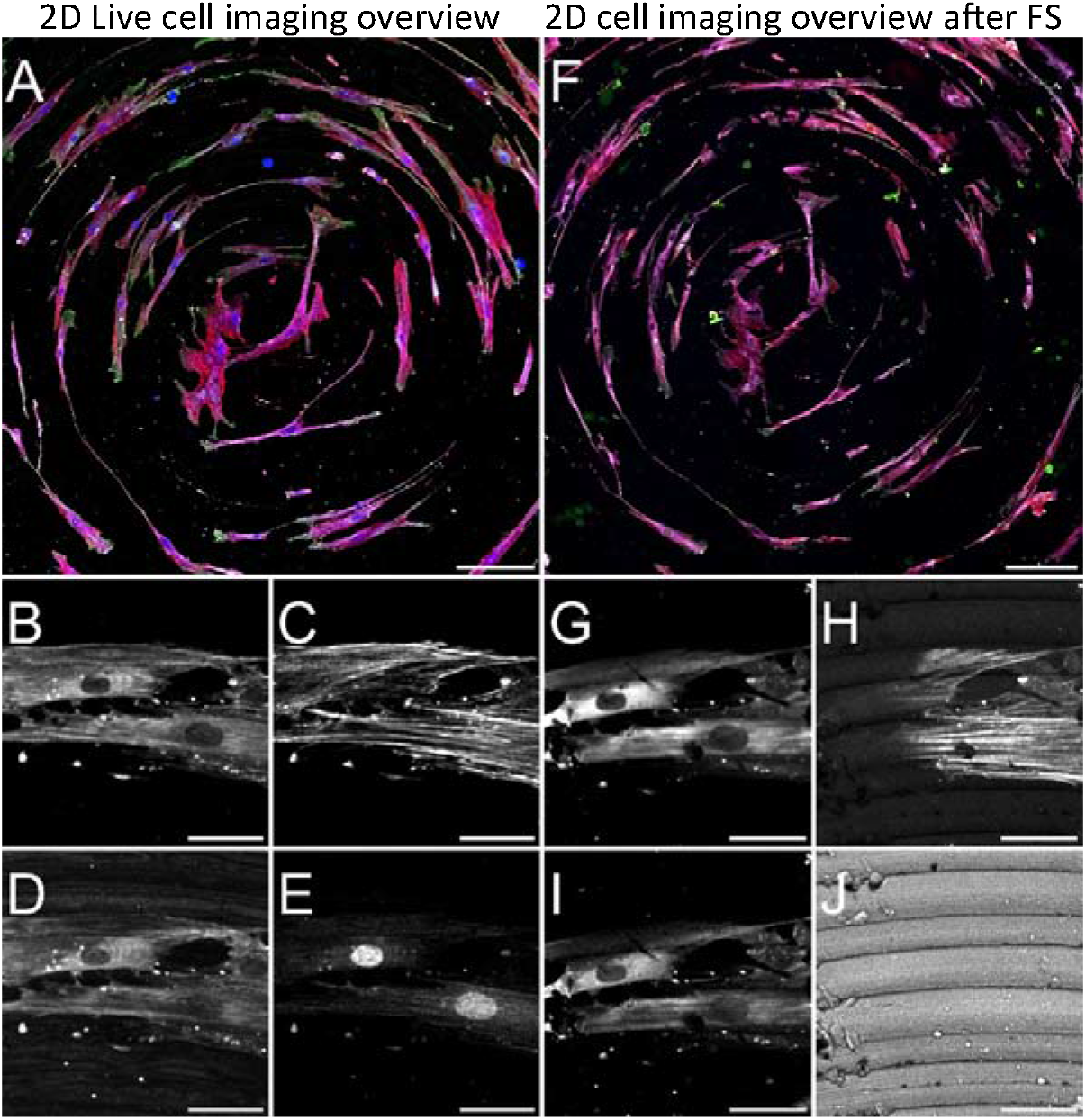
Fluorescence maintained in R221 resin after high pressure freezing and freeze substitution. Human osteoblasts (NHOst) grown directly on untreated HPF carriers and fixed with 4% PFA and 0.1% GA. The cells were permeabilized, then an antibody staining was performed (A-E) targeting clathrin (Alexa Fluor 647 B,G) and Osteocalcin (Alexa Fluor 488 D,I) and additional chemical dyes were used to stain actin (Phalloidin, Alexa Fluor 568, C,H) and nucleus (DAPI E,J). Next, cells were imaged on a Zeiss LSM900 and frozen using HPM Live μ. Subsequently, the cells were embedded into R221 resin using freeze substitution. Pre-HPF (A-E) and post-HPF/FS (F-J) images show the same areas of interest in both conditions. The Alexa Fluor dyes keep their fluorescence after HPF and FS, however DAPI fluorescence is lost after HPF and FS. Scale bars A,F 200 μm, B-J 50 μm

### iii. Correlating Fluorescence with SEM: towards 3D CLEM

Before transferring the sample to the SEM, the surface of the block should be smoothened by removing the first layer of resin with microtomy. The exposure of the surface leads to an increase of details in LM (Fig. 16). However, microtomy may remove valuable pieces of the sample (e.g. part of the cells), which can lead to loss of crucial information for 3D CLEM.

**Figure 16.**
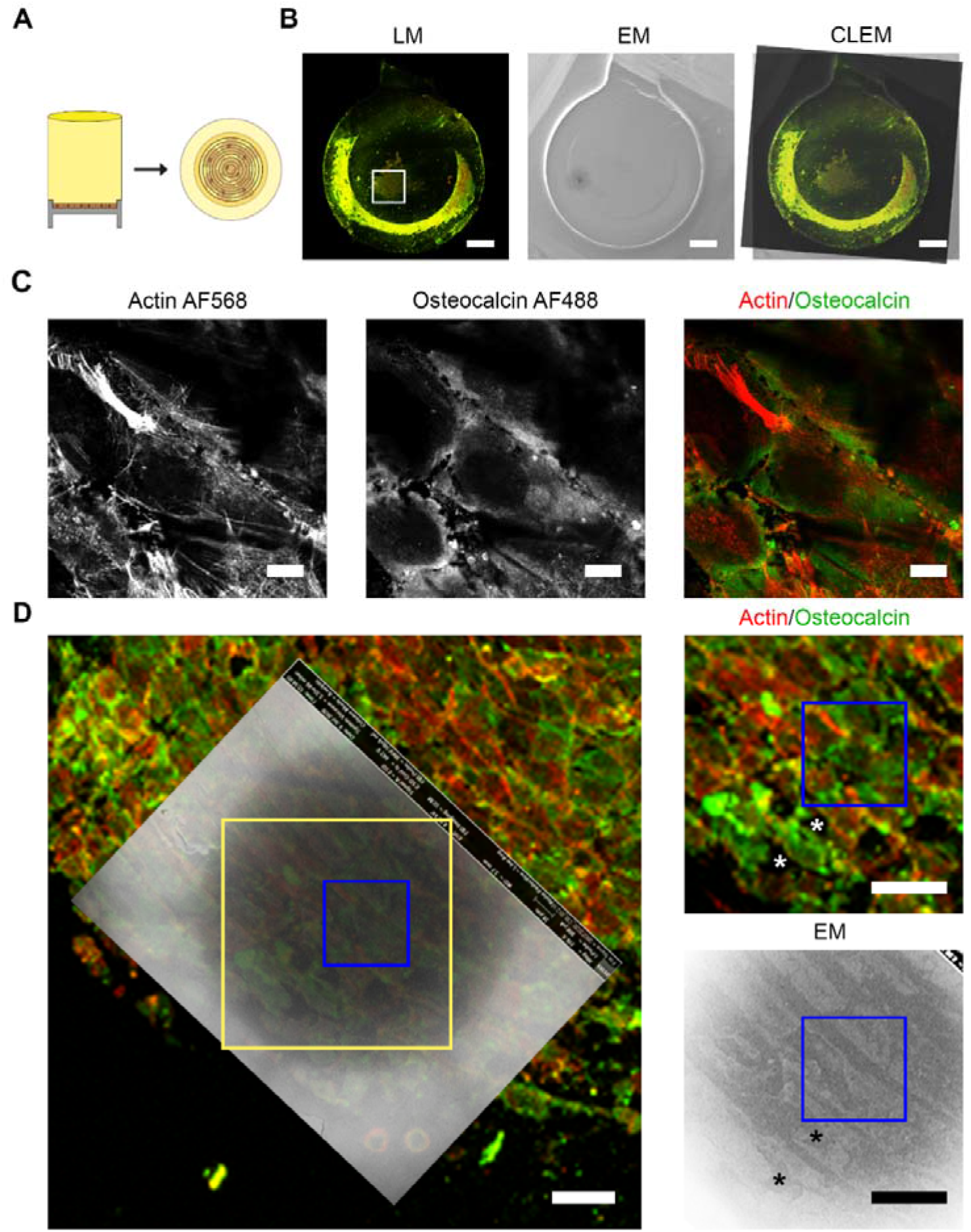
Correlation of fluorescence and scanning electron microscopy. (A) Cartoon representing the set-up. Cells grown on the carrier are embedded in plastic. Once the carrier is removed, the cells are at the top of the block. (B-D) MLO-A5 cells grown directly on the HPF carrier. Cells are fixed with 2% PFA and 0.1% GA, and subsequently permeabilized using 0.1% X-100 Triton. Cells are stained with anti-osteocalcin antibody (2^nd^ AB alexa fluor 488 nm) and phalloidin-Alexa Fluor 568 nm to stain the actin filament. Afterwards, the cells are frozen using HPF and embedded in R221 via freeze substitution. (B) The top of the block imaged with fluorescence light microscopy (LM) and by secondary electron SEM. The correlation of the two overview was used as a first rough CLEM alignment. White box: Area shown in D. Bars: 500 μm. (C) Region of interest imaged at high resolution. Pixel size: 0.043 x 0.043 x 0.150 μm. Here the image represents 8 Z slices. Note the preserved actin fibers. Bars: 10 μm (D) CLEM overlay was made using the backscattered electron SEM image. Blue box: high resolution LM images area shown in C. Yellow box: Zoomed in area at the right in D. * represents two of the anchor points used to create the correlation. It must be noted that only the surface is imaged with EM, so not all the cell outlines are visible as seen in LM. Bars: 50 μm.

This problem may be overcome by first coating the carrier with hydrogel or collagen layer, creating a spacer between the carriers and the cells. When using this spacing layer, the first layer removed by ultramicrotomy will only contain the collagen and resin, while the cells stay intact. The ability to observe more details in both LM and EM will improve the correlation on which CLEM critically depends.

Although a first preliminary correlation of the cryo-fluorescence and SEM imaging can be made based on the visual observation of cell morphology and other surface markers, the use of dedicated software is required to precisely locate the region of interest to be correlated (Paul-Gilloteaux et al., 2017). In addition, the cryo-fluorescence image will help in the alignment of the live images and the SEM image for post-acquisition analysis. Thus, monitoring each step contributes to the quality control of the experiments and thereby improving the efficiency of the imaging workflow and analysis with EM.

## 6. Perspectives

In this chapter we have shown how HPF in general and the HPM live μ in particular are at the crossroad of multimodal imaging, being a central player in sample preparation and preservation that is crucial to CLEM, and compatible with all contemporary CLEM approaches. Combining this unique HPM Live μ setup, the CryoCapsule and innovative embedding resin (R221^®^, CryoCapCell, France), the complete 3D Live cell to 3D EM CLEM workflow becomes achievable, imaging the sample by fluorescence at every step of the process. However some challenges still exist.

Recent years have shown great dynamics in the field of correlative microscopy, and a growing need in various light microscopy methods (live, fixed, cryo) to observe biological phenomena. While designing our new HPM live μ, the leading concept was to directly connect live imaging to the vitrification process. The first idea was to build an HPM with integrated confocal microscope. This would have had two benefits: it would potentially reduce of the total cost of the system (HPM + Microscope) and also minimize the total size of the combined system. The latter is still a point of attention, as the HPM Live μ is a large system and requires a dedicated location to be used under optimal conditions. Yet, the downside of this strategy would be the reduced flexibility of the integrated microscope (predefined and difficult to keep up with the state of the art).

In recent years, HPF has taken a central role in sample preparation for multimodal imaging, increasing the importance of monitoring and assessing the quality of the frozen sample. HPF is often summarized as a simple technique with little attention paid to the precise but crucial information on the freezing conditions used. The critical parameters that influence the final outcome of the vitrification quality (a combination of HPF curve, sample preparation and sample carrier selection) are usually not stored for long term reference, even though they could greatly help us to optimize freezing and sample preservation.

For this reason we introduced the advance graphical user interface and high accuracy measurements to register the events in the HPF chamber during the HPF shot. We record and store parameters for each individual sample, in order to explore them and allowing to optimize sample preparation. On the long term, this cumulated information will represent a source of knowledge to the lab and the growing HPF community. We envision that a more precise tuning of the HPF parameters, together with the sample preparation, will help vitrify samples that, until now, have been impossible to cryo-immobilize.

From an experimental perspective, the increasing variety of samples that require optimal ultrastructure preservation, will likely lead to the development of more complex sample carriers, allowing interaction with the sample while observing it prior to HPF. The limited functionality of the aluminum carriers forces us to think about designing new HPF carriers, with the CryoCapsule being the first of this kind.

The development of advanced optical microscopy since the mid-1990s and the recent renewed interest in electron microscopy, using cryo-EM, are opening up a new era for CLEM. It highlights the advanced sample preparation techniques already established in this field and encourages the development of new ones. In this context, we expect an increasing interest in vitrification by the light microscopy field, and a growing demand for flexible and well-controlled high-pressure freezing, approached both inside and outside the field of electron microscopy.

## 7. Acknowledgments

This work is the result of many years of collaboration. Among the main support received we thank ABRA Fluid AG, Widnau, especially Markus Frei for his technical support in developing the core of the HPM Live μ, we greatly acknowledge the Nikon Imaging Centre@Institut Curie, especially Lucie Sengmanivong. -CNRS and the Cell and Tissue Imaging (PICT-IBiSA), Institut Curie, member of the French National Research Infractucture France-BioImaging (ANR10-INBS-04).Lucy Collinson and the EM STP group at the Francis Crick Institute for supportive discussions and comments to improve the user experience on the HPM and the various engineers at Carl Zeiss that supported the technical integration of the Airyscan system. We would also like to thank Katarina Wolf for providing the fibrosarcoma cells. MdB, NS and AA were supported by the European Research Council (ERC) Advanced Investigator grant (H2020-ERC-2017-ADV-788982-COLMIN), AA is also supported by a VENI grant of the Netherlands Scientific Organization NWO (VI.Veni.192.094)

## 8. Author contributions

XH, JH, MB, LM, FE conceived and produced the HPM Live μ. XH, MdB, NS and AA conducted the proof of principle experiment. MKtL provided the cells for the experiments and conducted ultramicrotomy sectioning.

FS, JL, EL participated in the microscope integration and various associated tests of compatibility and stability. JS, GR supported the initial integration work at Institute Curie and provided technical and scientific resources. XH, MdB, NS, and AA wrote this chapter with the comments and additions of all authors.

## Notes

### Competing Interest Statement

The authors have declared no competing interest.

